# Unraveling the Genomic and Environmental Diversity of the Ubiquitous *Solirubrobacter*

**DOI:** 10.1101/2023.10.31.564804

**Authors:** Angélica Jara-Servín, Gerardo Mejia, Miguel F. Romero, Mariana Peimbert, Luis D. Alcaraz

## Abstract

2.

*Solirubrobacter*, a genus within the Actinobacteriota phylum, is commonly found in soils and rhizospheres yet remains unexplored despite its widespread presence and diversity, as revealed through metagenomic studies. Previously recognized as a prevalent soil bacterium, our study delved into phylogenomics, pangenomics, environmental diversity, and bacterial interactions of *Solirubrobacter*. Analyzing the limited genomic sequences available for this genus, we uncovered a pangenome consisting of 19,645 protein families, with 2,644 constituting a strict core genome. While reported isolates do not exhibit motility, we intriguingly discovered the presence of flagellin genes, albeit with an incomplete flagellum assembly pathway. Our examination of 16S ribosomal genes unveiled a considerable diversity (3,166 operational taxonomic units OTUs) of *Solirubrobacter* in Mexican soils, and co-occurrence network analysis indicated its extensive connectivity with other bacterial taxa. Through phylogenomic analysis, we delved into the relatedness of sequenced strains and notably dismissed ASM999324v1 as a member of this genus. Our investigation extended to the metagenomic diversity of *Solirubrobacter* across various environments, affirming its pervasive presence in rhizospheres and certain soils. This broader pangenomic view revealed genes linked to transcription, signal transduction, defense mechanisms, and carbohydrate metabolism, highlighting *Solirubrobacter’s* adaptability. We observed that *Solirubrobacter’s* prevalence in rhizospheres is geographically indiscriminate, prompting intriguing questions about its potential interactions with plants and the biotic and abiotic determinants of its soil occurrence. Given its richness and diversity, *Solirubrobacter* might be a versatile yet overlooked keystone species in its environments, meriting further recognition and study.

**Impact statement:** This study explored the enigmatic world of *Solirubrobacter*, a widespread microbe commonly found in soils and plants across various regions. Despite its prevalence, little is known about its genetic diversity and functionality and how it thrives in diverse environments. Our research unveils the genetic secrets of *Solirubrobacter*, shedding light on its adaptability and ecological interactors and roles. We showed that *Solirubrobacter* environmental prevalence makes it a good candidate for studying the genetic basis of being a successful microbe associated with soil and plants.

**Data summary:** Data, scripts and statistical analysis available in GitHub: https://github.com/genomica-fciencias-unam/Solirubrobacter

Sequences, phylogenetic analysis, raw data structures: https://doi.org/10.6084/m9.figshare.24446521

16S rRNA gene raw data: https://www.ncbi.nlm.nih.gov/sra/PRJNA603586

https://www.ncbi.nlm.nih.gov/sra/PRJNA603590

Shotgun metagenomes: https://www.ncbi.nlm.nih.gov/bioproject/603603

All supporting data, code, and protocols are within the article, supplementary files, and described repositories.

## 5. Introduction

*Solirubrobacter* is an Actinobacteriota bacterial genus whose abundance appears high in soils worldwide [1]. The genus’s first type strain and definition was isolated from vermicompost in 2003, *Solirubrobacter pauli* [2]. *S. pauli* was described as a genus of Gram-positive bacilli, pink-colored, aerobic and mesophilic, non-sporulating, without motile structures, low desiccation resistance, no growth at 1% NaCl, and with a ∼70% G+C content [1,2]. Since then, *S. soli*, *S. ginsenosidimutans, S. phytolacca,* and *S. taibaiensis* have been isolated and described. Both *S. soli* and *S. ginsenosidimutans* were isolated from ginseng fields, while *S. phytolacca* and *S. taibaiensis* from the roots and stem of the Indian pokeweed *Phytolacca acinosa* [3–5]. The carbohydrate utilization capabilities, growing conditions, and phenotype vary depending on the species (Table 1). The cultivated isolates of *Solirubrobacter* are very limited, probably due to their slow growth and the lack of a specific selective medium, even though poor nutrient media seem to contribute to their isolation [6]. *Solirubrobacter* is also related to harsh environments, such as black chickpea fertilized culture soils [7] and coal and soil samples from opencast coal mines [8]. This bacterial genus has been linked with metabolizing contaminants such as detergents [9] and positively correlated with bioaccumulation of Cd and Zn in plants [10]. Despite being widely reported in agricultural soils, *Solirubrobacter* seems sensitive to antibiotics, decreasing its abundance by more than 50% in agricultural soils contaminated with antibiotics of manure origin [11]. Even though some strains have been reported as non-resistant to desiccation [2], in studies analyzing UV exposure, *Solirubrobacter* appeared in slightly increased abundance [12] and even as one of the key genera in microbial communities from sand desert exposed to intense solar UV radiation [13].

**Table 1.** *Solirubrobacter* reference genome information and traits.

| Strain | Accession | Size (Mb) | %GC | Completeness | Country | Color | Growth T (°C) | Growth >1% NaCl | Motility | Sporulation | Environment | Genome submitter | Associated publications |
| --- | --- | --- | --- | --- | --- | --- | --- | --- | --- | --- | --- | --- | --- |
| *S. soli* DSM22325 | ASM42366v1 | 9.31 | 71.4 | 98.71 | South Korea | White | 15-33 | - | - | - | Ginseng field | DOE Joint Genome Institute | Kim et al., 2007, Mukherjee et al., 2017 |
| *S. sp* URHD0082 | ASM42594v1 | 6.64 | 72.3 | 99.14 | N.d. | N.d. | N.d. | N.d. | N.d. | N.d. | Mediterranean grassland | DOE Joint Genome Institute | N.A. |
| *S. sp* CPCC204708 | ASM334462v2 | 7.59 | 69.9 | 98.71 | China | N.d. | N.d. | N.d. | N.d. | N.d. | Desert sand | Chinese Academy of Medical Sciences & Peking Union Medical College | N.A. |
| *S. pauli* | ASM363375v1 | 7.13 | 72.1 | 98.71 | USA | Pink | 28-30 | - | - | - | Vermicompost | DOE Joint Genome Institute | Singleton et al., 2003 |
| *S. ginsenosidimutans* | ASM2758720v1 | 9.69 | 71 | 99.57 | China | Pale yellow | 18-37 | - | - | - | Ginseng field | Chinese Academy of Medical Sciences & Peking Union Medical College | An et al., 2011 |
| *S. phytolaccae* | ASM2758719v1 | 7.73 | 71.5 | 98.71 | China | White | 25-30 | + | - | - | Root of *Phytolacca acinosa* | Chinese Academy of Medical Sciences & Peking Union Medical College | Wei et al., 2014 |

Numerous environmental studies have detected this bacterium through 16S rRNA gene sequencing, especially in soils and rhizospheres of diverse plants. Our group has examined soil and rhizosphere microbiome samples across Mexico, from soils with very diverse physicochemical properties and rhizospheres of wild and cultivated plants. In these hundreds of samples we have observed the prevalence of *Solirubrobacter* [14–16]. The limited data on this genus obstructs elucidating the precise role *Solirubrobacter* plays in the rhizosphere. However, both favorable and adverse correlations with plant growth have been documented [15, 17, 18], implying potential gene presence promoting host plant development in *Solirbrobacter* genomes. Interestingly, *S. soli* DSM 22325 was sequenced as a directed effort of the Genome Encyclopedia of Bacteria and Archaea (GEBA) [19]. The GEBA consortium also identified *S. soli* as the top recruiting genome when aligning it against metagenomes, with preferences for terrestrial (50%), plant (34%), and aquatic environments (6.5%) [19].

The widespread prevalence of this bacterial genus resonates with the assertion, “Everything is everywhere, but the environment selects,” a notion sparking ecological debates since the eighteenth century [20]. This theory posits that due to the swift population expansion and vast size of microbial communities, microbial taxa can disperse rapidly, and their distributions are shaped by environmental factors—thus negating the roles of historical biogeography and dispersal events in community formation [21, 22]. Among these environmental determinants are established factors like pH, alongside more intricate ones such as pedogenesis, trophic status, available plant debris, and vegetative cover [23, 24]. However, some research indicates that environmental influences wane beyond a specific geographic extent (10 km), which suggests that the same biogeographic and historical processes of macroorganisms can apply to bacterial populations and communities [21, 25]. Geographical barriers foster endemic microorganisms, primarily as they prevent a significant dispersion [21, 26]. Even so, instances of microorganisms from the same taxonomic group identified across disparate geographical locations have been recorded, with the underexplored genus *Solirubrobacter* being one such case. Despite its ubiquity across diverse soils and plant hosts, the genetic diversity enabling *Solirubrobacter* to flourish in varied and contrasting environments remains overlooked. Delving into the genetic makeup of *Solirubrobacter*, in light of the “Everything is everywhere, but the environment selects’’ concept, not only elucidates this genus but also sheds light on how geographically distant locales and differing environmental attributes might influence the genetic distribution and adaptive strategies of a singular bacterial genus.

*Solirubrobacter* is a common and prevalent bacterium in soils and rhizospheres, yet its biology remains largely unexplored. This study aims to unravel the diversity and functionality of *Solirubrobacter*, striving to elucidate its ecological roles, adaptive capabilities, and interactions with other microorganisms. To achieve this, we adopted a comprehensive strategy encompassing phylogenomics, pangenomics, and metagenomic analysis. Initially, we collected and examined the available genomic sequences of the *Solirubrobacter* genus. Additionally, we analyzed environmental 16S ribosomal gene sequences to discern the diversity and distribution of *Solirubrobacter* across various environments. Subsequently, we constructed an Environmental Extended Pangenome (EEP) by recruiting potential new *Solirubrobacter* proteins from soil and rhizospheric metagenomes to investigate if similar environments select for comparable features. These methodologies enabled us to determine the prevalence of *Solirubrobacter* in soil and rhizospheres, regardless of geographic location. Our integrated approach enhanced the understanding of *Solirubrobacter* genomic diversity and its significance in terrestrial ecosystems, laying a foundation for further investigation and continuous study of this understudied microbial genus.

## 6. Methods

### *Solirubrobacter* core genome retrieval and phylogenetic analysis

We summarized the main methodological steps (Fig. 1). All available genomes and draft assemblies from the genus *Solirubrobacter* were retrieved from NCBI in February 2023. In total, eight *Solirubrobacter* species were downloaded (Table 1). Genome quality and completeness were evaluated using Checkm [27]. Using the GET_HOMOLOGUES pipeline [28], we calculated a pangenome of all *Solirubrobacter* strains while using *Streptomyces coelicolor* and *Bacillus subtilis* as outgroups (NCBI Accessions: GCA_013317105.1 and GCF_000009045.1). Core genes were retrieved and aligned individually with MAFFT [29] for further concatenation into a core genome alignment. Then, using R v 4.2.2 [30] ape [31] and phangorn [32] libraries, we calculated the best evolutionary model of the core genome alignment with ten taxa, a length of 105,813 characters, and 48,460 site patterns. The best Akaike information criterion (AIC) model was LG [33], with a gamma-distributed rate (k = 4). The model was used to build a consensus maximum likelihood phylogeny with 1,000 bootstraps. The complete analysis protocol is available on GitHub (https://github.com/genomica-fciencias-unam/Solirubrobacter). Additionally, we carried out the genome similarity score (GSS) analysis [34–36]. A set of seven *Solirubrobacter* genomes was retained and used for further analysis and are henceforth called reference genomes or reference species.

**Fig. 1.**
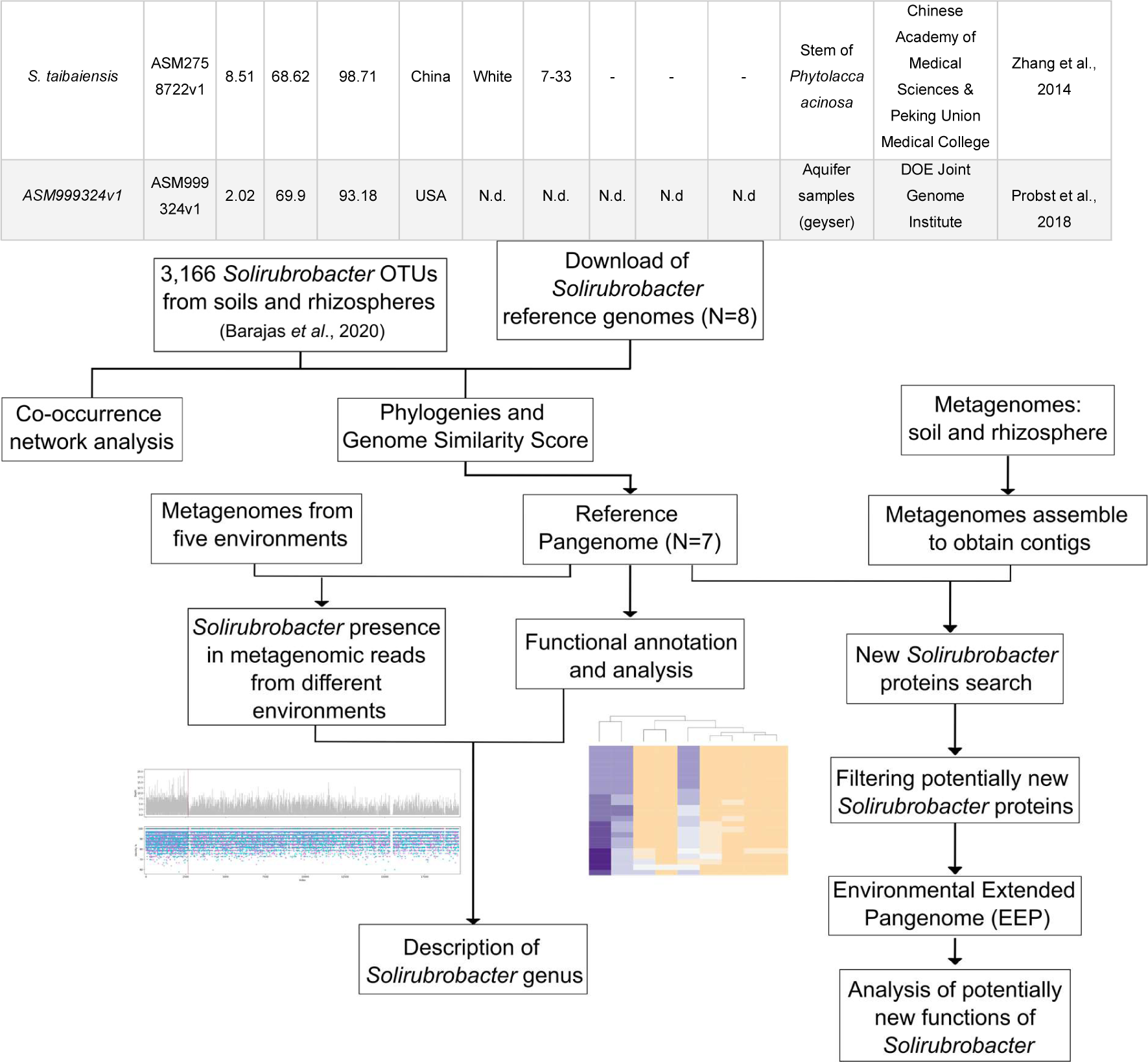
Overview of this work.

### *Solirubrobacter* diversity and networks using 16S rRNA gene

We reanalyzed 107 microbiomes of rhizosphere, endosphere, and soils from Mexico [14] to extract the sequences of the OTUs corresponding to *Solirubrobacter*. The dataset included 255,275 16S OTUs (97% identity). For genus-level analysis, the dataset was managed with R’s phyloseq [37], vegan [38], and ggplot2 [39] libraries. We also extracted the 16S rRNA genes from the reference genome sequences using barrnap (v. 0.7) [40] and concatenated them into the 16S microbiome sequences. We aligned all the 16S using SSU-align [41], followed by a phylogeny using FastTreeMP [42].

For the network analysis, the 16S OTUs sequences were merged to the genus level, resulting in 1,748 bacteria genera. With the R igraph library [43], we calculated a co-occurrence, undirected network. We exported it as a graphml object to be analyzed by Gephi graph visualization and manipulation software (v 0.10) [44]. Complete protocols and datasets are available on GitHub (https://github.com/genomica-fciencias-unam/Solirubrobacter) and FigShare (https://doi.org/10.6084/m9.figshare.2444652).

### *Solirubrobacter* pangenome analysis

All *Solirubrobacter* genomes were annotated using Prokka (v.1.12) [45] to generate gff3 output files that served as input for Roary [46]. Roary is designed to construct pangenomes from related species. An UpSet diagram [47] was constructed with R to illustrate the shared and unique gene distribution. The resulting output files with the sequences comprising the pangenome were annotated using the Clusters of Orthologous Groups (COG) [48] and Kyoto Encyclopaedia of Genes and Genomes (KEGG) [49] database. The resulting annotations were used to create a matrix with counts of the number of proteins of each COG in the reference pangenome, the core genome, and each *Solirubrobacter* strain. The matrix was standardized using Z values. Using only the sequences successfully annotated in COG, a functional profile was constructed using the phyloseq, ggplot2, gplots [50], and R default packages. Hierarchical clustering was performed using the *hclust* method on a Bray-Curtis dissimilarity matrix [51].

After annotation, we were able to detect proteins involved with flagellar biosynthesis. Considering that *Solirubrobacter* isolates are described as non-motile, we retrieved all proteins involved with flagellar assembly (KEGG and COG databases). All selected proteins were used to construct a presence-absence matrix. For comparison, the reference genomes of *Conexibacter woesi* and *Patulibacter minatonensis* were downloaded from NCBI (Accession number: GCA_013317105.1 and ASM51932v1, respectively) and annotated against the same databases. We chose these bacteria because they are flagellated and belong to the same order as *Solirubrobacter*.

### *Solirubrobacter* recruitment with metagenomes from multiple environments

To evaluate the presence of the reported *Solirubrobacter* species in environmental samples, we used a total of 36 metagenomes from rhizospheres and soils from different geographic locations. Mexican soils and rhizospheres comprised 16 of the 36 selected metagenomes and were retrieved from a previous study evaluating soil and rhizosphere microbial composition (NCBI Bioproject PRJNA603603) [14]. The remaining soil metagenomes were selected if the sample came from an agricultural field and the location was geographically distant from our Mexican soils. The criteria for rhizospheres was also a geographical separation and plant host different from the Mexican ones. Lake water, phyllosphere, and sea sediment metagenomes were used as negative controls. All metagenomes used for this section are listed in Table S1. Reads associated with *Solirubrobacter* from the 36 metagenomes were identified using the reference pangenome sequences and tBLASTn [52]. The resulting hits were filtered according to bitscore value and quality-checked using Trimmomatic [53]. Later, the constructed pangenome and the *Solirubrobacter* metagenomic reads were aligned using Promer [54]. We then generated identity graphs showing the percentage identity and coverage of the alignment through the *promer_deid.py* in-house script available on GitHub. After analysis of the Promer alignments, the rest of the study was carried out using only the metagenomes from Mexican soils (six metagenomes. Samples 1 to 6; Table S1) and Mexican rhizospheres (ten metagenomes. Samples 13 to 22) were selected for the construction of the Environmental Extended Pangenome (EEP).

### Metagenomic data assembly

The sixteen selected Mexican soils and rhizosphere shotgun metagenomes from the NCBI Bioproject PRJNA603603 [14] were subjected to quality control using Trimmomatic, and paired-end matched reads were retained for subsequent analysis. Metagenomic reads from samples 18 to 22 (rhizospheres; Table S1) were filtered out, matching *Solanum lycopersicum* genome (Accession number: GCF_000188115.5), while samples 13 to 17 (rhizospheres; Table S1) were filtered against the *Arabidopsis thaliana* genome (Accession number: GCF_000001735.4) with Bowtie2 [55]. Metagenomic libraries were assembled using a hybrid assembly approach. First, SPADES [56] with –a meta function was used to generate contigs, and high-quality reads were mapped against the SPADES contigs. Second, all unmapped reads were subjected to a second assembly with Velvet [57]. The resulting contigs from both assemblies were merged, and only a minimum of 100 bp length contigs were retained. All obtained contigs were used to obtain an EEP for *Solirubrobacter*.

### *Solirubrobacter* Environmental Extended Pangenome

To recruit potentially new *Solirubrobacter* proteins, we used the pangenome sequences as a reference and, using Bbmap [58], mapped them against the previously assembled contigs of the 16 selected metagenomes. All contigs containing *Solirubrobacter* proteins were extracted, and coded proteins surrounding the already known *Solirubrobacter* protein were considered potentially new proteins of this genus. We performed a filtering step using BLASTx [59] with the pangenome as a database and filtering by an e-value of 1e-5. We then used Prodigal [60] to predict ORFs and coding proteins for the new sequences and the reference sequences of *Solirubrobacter*. All sequences were concatenated in a single file and ran through CD-HIT [61] using the following parameters: -n 3 -c 0.50 -aS 0.7 -d 0 -M 30000 -T 10. These settings allowed us to create an EEP composed of the potentially new *Solirubrobacter* proteins and the proteins from the reference species. The EEP was annotated using COG and KEGG as carried out for the reference pangenome.

### Functional and diversity analysis of the *Solirubrobacter* environmental extended pangenome (EEP)

Using only the sequences successfully annotated in COG, a functional profile of the EEP was constructed using the same methods listed for the reference pangenome functional profile. Additionally, for the annotated sequences, differences between samples were analyzed through an unconstrained Principal Coordinate Analysis (PcoA) [62] on Bray-Curtis dissimilarity matrices; the clustering was evaluated through the ANOSIM statistical function [63]. To further describe the metabolic capability of the genus *Solirubrobacter*, the number of duplicated sequences per sample was counted using the output file of the CD-HIT analysis. This analysis allowed us to determine gene families and evaluate which metabolic activities *Solirubrobacter* assigns a higher portion of its genome. Detailed statistical and bioinformatic methods for all the methods used in this study can be accessed on GitHub.

Annotated sequences from the reference and EEP were manually analyzed to identify any proteins related to plant-microorganism interactions. Additionally, Hidden Markov Models (HMM) for Actinobacteria proteins involved in plant-microorganism interactions [64] were used to search for proteins of this type of interaction in *Solirubrobacter* metagenomic proteins from Mexican soils and rhizospheres. This search used HMMER v. 3.3.2 [65], and only the three best hits were requested. *Streptomyces coelicolor* and *Streptomyces griseus* genomes (Accession number: GCA_013317105.1 and ASM1060v1, respectively) were used for comparison since strains from these species are known to interact with plants.

## 7. Results

### Functional composition of *Solirubrobacter* references core genome

A total of 13,160 proteins were annotated (66.98% of the pangenome, Table S3) using the COG ontology, with 2,421 of them belonging to the core genome. Hypothetical proteins accounted for 6,484 of the pangenome, of which 223 were from the core genome. We visualized the abundance of the COGs per species, as well as the core genome and pangenome (Fig. 2). The most abundant COGs in the pangenome were S (unknown; 18.19%), K (transcription; 11.53%), T (transduction; 6.44%), E (aminoacids; 6.29%), and G (carbohydrate; 5.94%). Regarding annotated sequences from the core genome, the most abundant COGs were S (unknown; 15.61%), E (aminoacids; 8.01%), K (transcription; 7.47%), C (energy; 6.81%), NA(-)(6.07%), J (ribosomal; 5.86%), and G (carbohydrate; 5.24%).

**Fig. 2.**
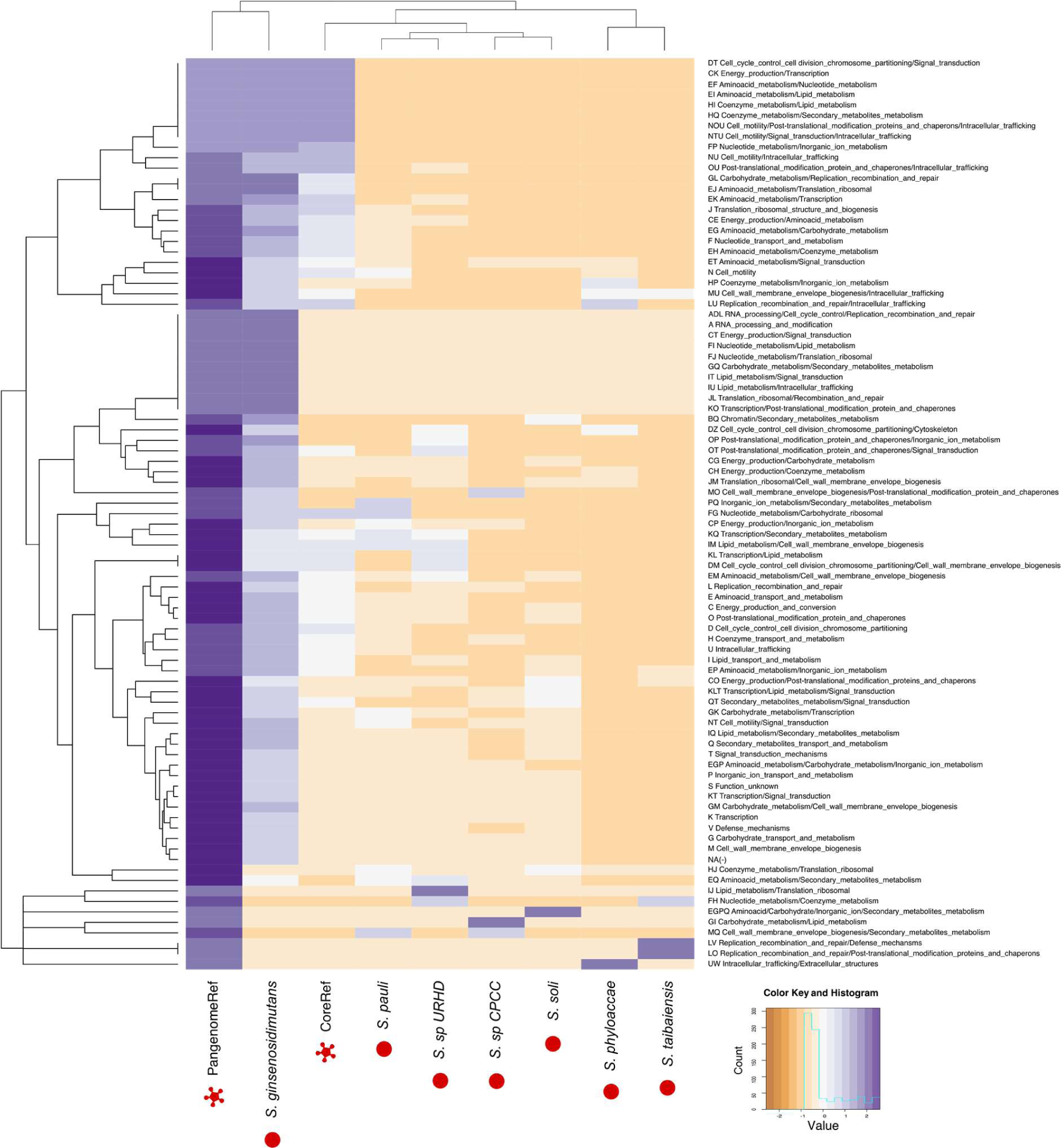
Functional profile of *Solirubrobacter* references pangenome. Heatmap showing normalized Cluster of Orthologous Groups (COGs) frequencies within each group are shown as a two way clustering. Circles refer to reference strains and stars to the constructed pangenome and core genome of *Solirubrobacter*.

The ability to exploit different carbohydrate sources is reflected in the core COG G (carbohydrate). In this COG we found five protein families identified as ROK family proteins, as well as enzymes for xylose isomerases (8), alpha amylase domains (2), starch synthase (1), hydrolases (12), HpcH/HpaI aldolase/citrate lyase family (1), and polysaccharide deacetylase (2). Moreover, a total of 8 cupin proteins or cupin domains were identified between COGs G and S (carbohydrate and unknown, respectively).

Sigma factors lead to a wide gene expression diversity by modifying the specificity of RNA polymerase [66]. Among COGs K (transcription), T (transduction), and KT (transcription/transduction) of the core genome we detected 25 sigma factors, of which three were ECF sigma factors and 13 sigma-70 factors. Histidine kinases are an important part of two-component systems environmental sensing [67] and a total of 18 annotated protein families were histidine kinases. Additionally, 12 protein families were identified as the helix_turn_helix domain of the Lux regulon or as LuxR regulators.

Regarding defense mechanisms (COG V), ABC transporters represented 26.47%, including daunorubicin resistance (4) and chromate (1) transporters. We detected homologs to genes *drrA* and *drrB*, conferring resistance to daunorubicin, an antibiotic produced by bacteria from the genus *Streptomyces* [68, 69]. Two family proteins were annotated as fungal trichothecene efflux pump (TRI12), contributing to the secretion of this mycotoxin from the cell. The AAA domain from the putative AbiEii Type IV Toxin-Antitoxin system was also identified in the core genome. AbiE TA systems confer phage resistance to the cell-inducing bacteriostasis: AbiEii acts as a GTP-binding NTase toxin that is neutralized by the expression of AbiEi [70]. A total of 18 protein families were annotated as dioxygenases, including glyoxalase/bleomycin resistance and protocatechuate dioxygenases, key enzymes in the degradation of various aromatic compounds [71]. ArsB, an arsenic pump membrane along cognate regulators, was identified in the core genome. Stress-related proteins were detected. Three families of cold shock proteins are involved in cellular responses against low-temperature stress, while four protein families are involved in the use of trehalose as a compatible solute (COG G; carbohydrate). The protein YaaA is involved in peroxide stress (COG S; unknown). From COG P (inorganic ions), the iron-storage gene *bfrB* gene was also found in the core genome. Finally, multiple repair systems for damaged DNA were found in 20 family proteins from the COG L (replication and repair), including the *uvrABC* operon. Secondary metabolism (COG Q) in the core genome is just 13.79%, while 86.21% of detected genes were in the accessory genome. In the core genome, three protein families were identified as cytochrome P450, while ten were identified as methyltransferases involved with secondary metabolism.

Upon annotating the core genome using KEGG, 52.8% of the core sequences were successfully annotated, enabling the reconstruction of biosynthetic pathways for tryptophan, phenylalanine, tyrosine, histidine, lysine, and the branched-chain amino acids valine, leucine, and isoleucine (Fig. S1). The metabolic pathway for arginine degradation to ornithine was identified, contrasting with the absence of an arginine biosynthetic pathway. The reference core genome lacked enzymes for nitrate reduction, yet enzymes and an ammonium transporter necessary for glutamate synthesis from ammonia were present. The *argFGH* operon for arginine biosynthesis was missing in the core (only *argH* was detected through COG annotation). Transporters for the missing amino acids were not identified in the core genome (except for the branched-chain amino acid transporter). However, a methionine transporter was found in the pangenome, along with portions of cystine and urea transporters. This suggests that for other essential amino acids, *Solirubrobacter* likely relies on membrane transporters for uptake.

### Gene families expansions in *Solirubrobacter*

We clustered all the proteins into single-copy genes, gene families from two to five members, and equal or larger than six within each *Solirubrobacter-*predicted proteome (Table S4). The most prominent gene families accounted for a maximum of 22 genes in *S. ginsenosidimutans* and *S.* sp. CPCC. They were annotated as COGs K (transcription) and L (replication and repair) (Fig. 3a). COG K (transcription) proteins were identified as helix_turn_helix Lux Regulon and COG L (replication and repair) as RNA-mediated transposition proteins. Gene families with nine genes coding for flagellin from COG N (motility) are present in strains *S. pauli, S. phytolaccae*, and *S. soli*. All strains, except for *S. phytolaccae* and *S.* sp. CPCC, have gene families from COG KT (transcription/transduction) of sizes ranging from seven to up to 15 and were identified as LuxR family transcriptional regulator genes. Interestingly, six families containing six or more hypothetical proteins were found. Of these hypothetical proteins, we analyzed family number 528 through a structural alignment that suggests that this family is homologous to the antimicrobial peptide Brevinin-1JDa (Fig. S2).

**Fig. 3.**
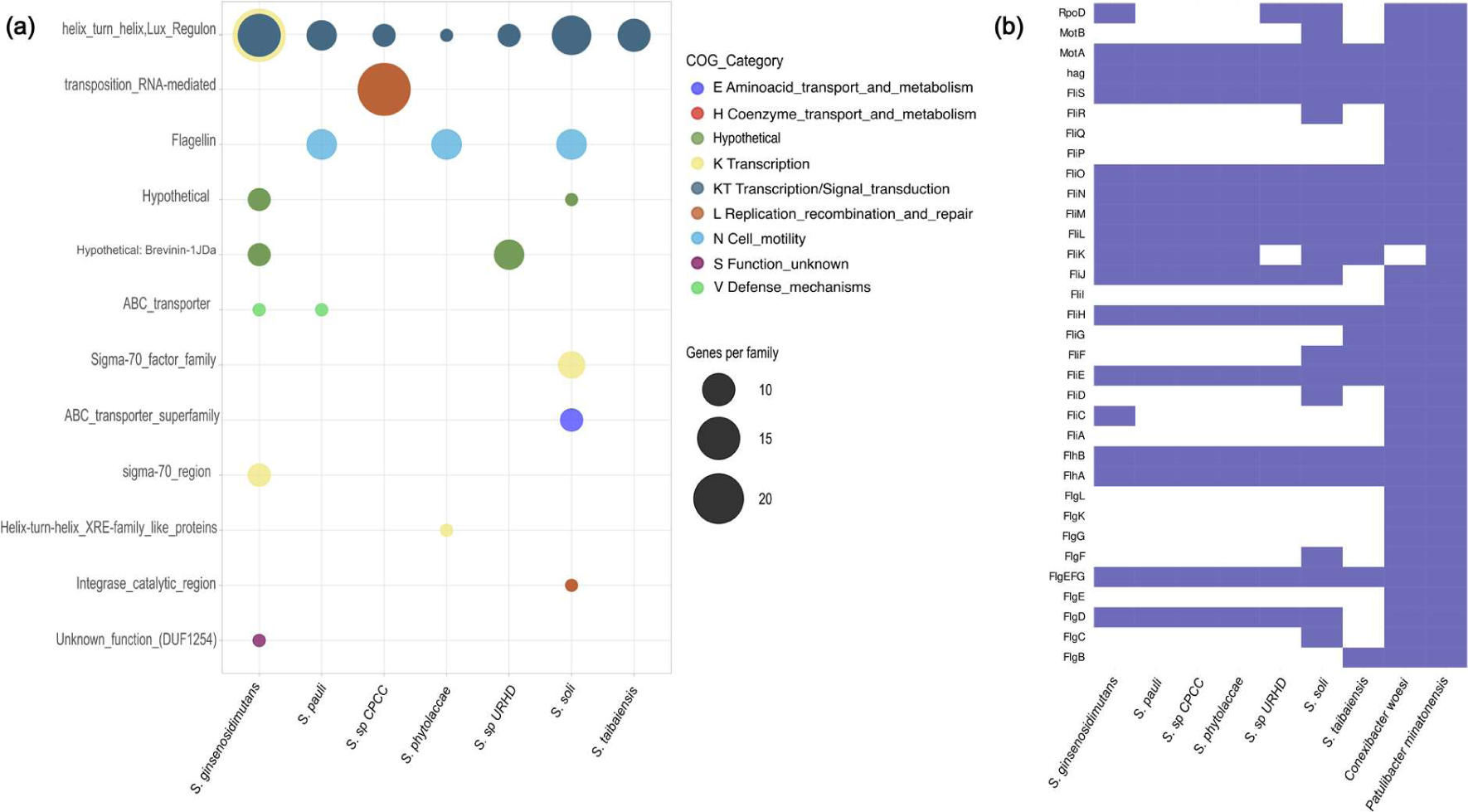
Extended families in *Solirubrobacter* references. (a) Bubble plot showing gene families with six or more duplicated genes and their annotation according to Cluster of Orthologous Groups (COGs) for each reference strain. (b) Detected proteins involved in flagellar biosynthesis in *Solirubrobacter*. The presence (blue) or absence (white) of each protein per *Solirubrobacter* reference is displayed. Flagellar biosynthesis proteins present in *Conexibacter woesi* and *Patulibacter minatonensis* are shown as comparison, because they are *Actinobacteriota* reported to swim. All flagellar proteins shown were selected based on the annotation using COG and KEGG databases.

Proteins involved with motility and flagellar assembly were detected among *Solirubrobacter* references (Fig. 3b). A total of 33 proteins bound to flagellar biosynthesis were identified in the Actinobacteriota flagellated species *Conexibacter woesi* or *Patulibacter minatonensis*, from which only 11 (33.33%) were present in all *Solirubrobacter* references. The rest of the proteins were detected in either *C. woesi* or *P. minatonensis* but were absent in most of the references. All *Solirubrobacter* and Actinobacteriota controls contained flagellin synthesis genes (*hag*). Moreover, gene families with nine genes coding for flagellin from COG N (motility) are present in strains *S. pauli, S. phytolaccae*, and *S. soli* (Fig. 3a). Among the genes detected in *Solirubrobacter* are structural components such as the hook (FlgE), along with regulatory elements like FliK, responsible for controlling hook size, and FliS, a chaperone facilitating flagellin transport [72]. Prime proteins such as the cytoplasmic membrane ring (FliF) and the motor component (MotB) are absent in *Solirubrobacter*.

### *Solirubrobacter* comparative genomics

We used the eight NCBI available genomes of *Solirubrobacter pauli* (type strain), *S. soli* DSM22325 (*S. soli*)*, S.* sp. CPCC204708 (*S.* sp. CPCC), *S.* sp. URHD0082 (*S.* sp. URHD)*, S. ginsenosidimutans, S. phytolaccae, S. taibaiensis,* and ASM999324v1. The main genomic features and reported phenotypic descriptors were summarized in Table 1. The genome size of ASM999324v1, annotated as a *Solirubrobacter* in GenBank, is considerably smaller (2.02 Mb) than the average genome size of the rest of the *Solirubrobacter* (8.08 ± 1.12 Mb). Using a 16S rRNA gene recovered from the genome sequences we did a phylogenetic reconstruction using *Streptomyces coelicolor* and *Bacillus subtilis* as outgroups (Fig. 4a). From the 16S of the genomes, the basal clades have bootstrap support (>0.6). In contrast, the top clade formed by *S. pauli* and *S. phytolaccae* had 0.5 support. We noted that ASM999324v1 is within the outgroup clade at all times (bootstrap = 1), with low support of the inner branching (Fig. 4a). Then, using a core genome maximum-likelihood phylogeny, we observed complete support (bootstrap = 1) in the consensus tree (1,000 bootstraps) for all branches of *Solirbubrobacter* and the outgroup clade by *B. subtilis* and *S. coelicolor* but with a single branch of ASM999324v1 (Fig. 4b).

**Fig. 4.**
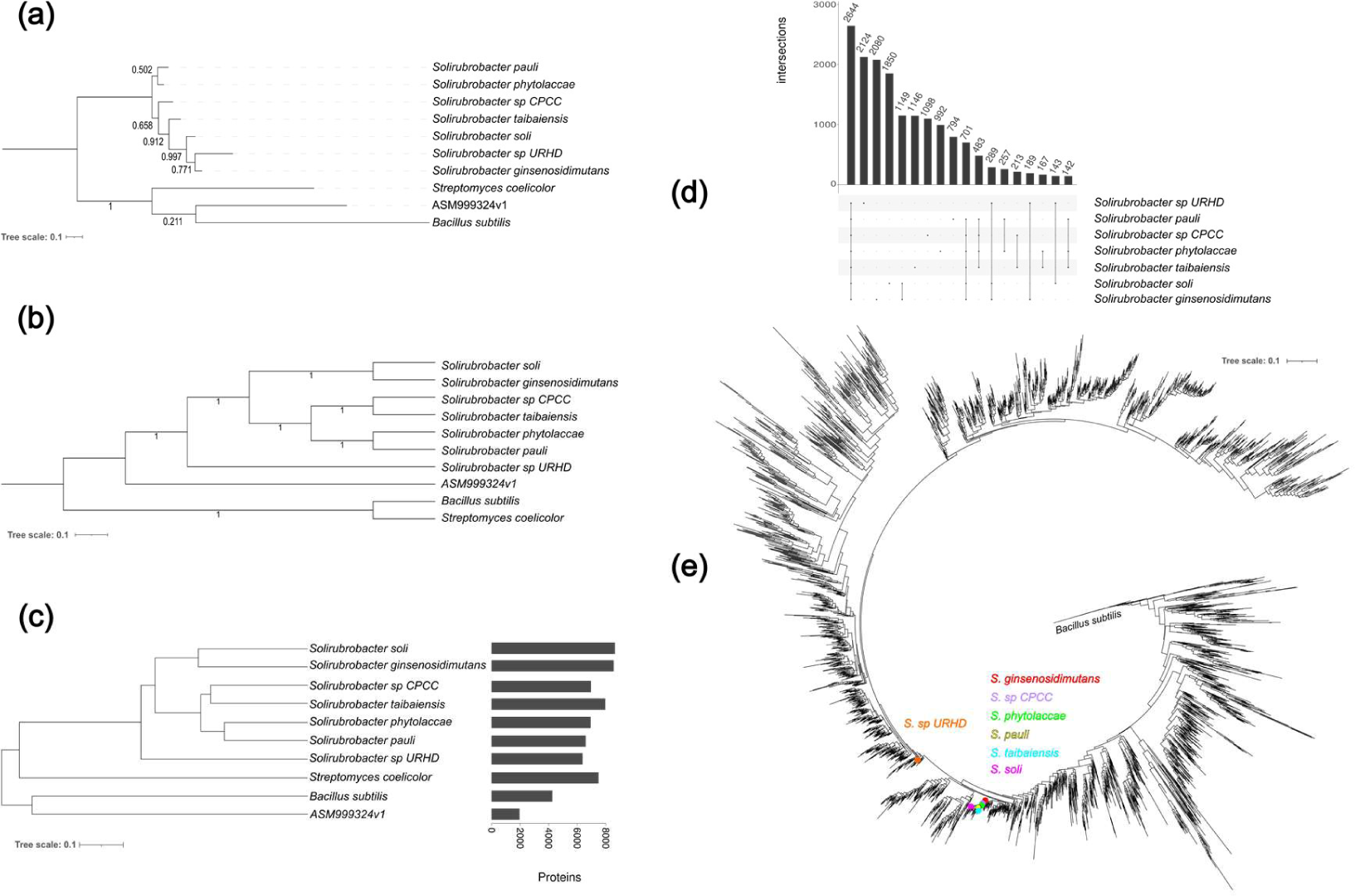
Phylogenomic diversity and shared genes of *Solirubrobacter*. (a) A 16S rRNA gene Neighbour-Joining (NJ) phylogeny of the available genome references. Bootstrap values are shown. (b) Core genome maximum likelihood phylogeny using 275 ortholog proteins and 1,000 bootstraps. (c) *Solirubrobacter* references dendrogram constructed using genomic similarity score (GSS) distance (d) UpSet diagram showing *Solirubrobacter* shared genes and pairwise intersections, in the first column, the core genome intersection is shown with 2,644 genes shared in all *Solirubrobacter* genomes. (e) *Solirubrobacter* reference genomes against environmental diversity, a 16S NJ phylogeny of the operational taxonomic units (OTUs) annotated as *Solirubrobacter* from Mexican soils and rhizospheres microbiomes, the reference genomes 16S are shown in colored dots. In all the phylogenetic and GSS analysis, *Streptomyces coelicolor* and *Bacillus subtilis* are used as outgroups.

The core genome consensus phylogeny showed *S.* sp. URHD as the basal clade, sister to two other clades, one formed by *S.* sp. CPCC, *S. taibaiensis, S. phytolaccae*, and *S. pauli*, and a top clade branched into *S. soli* and *S. gingsenosidimutans* (Fig. 2b). Both 16S and core genome phylogenies located ASM999324v1 in a separate branch than the rest of species from *Solirubrobacter*. Additionally, we conducted a genome similarity score (GSS) analysis because it is a metric that allows pairwise orthologs beyond the core genome, thus including vertically and horizontally transmitted elements and pangenomic information in the analysis. Briefly, the GSS used paired comparisons based on the sum of bit-scores of shared orthologs, detected as reciprocal best hits (RBH), and normalized against the sum of bit-scores of the compared genes against themselves (self-bit-scores). Its values range from 0 to 1, where 0 indicates the genomic content of compared species is the same, and a value of 1 indicates the absence of homologous proteins. Then, the GSS can be plotted as a distance dendrogram, where the maximum pairwise protein comparisons are plotted to the right of the graph (Fig. 4c). All *Solirubrobacter* strains had GSS values between 0.1896 - 0.3794, but ASM999324v1 presented an average value of 0.6615 when compared to the other strains of the genus (Table S2).

Additionally, the core genome and GSS provide valuable gene function information and allow us to identify shared functions and metabolic pathways among species, as well as provide information about the ecological roles and adaptations of the species. The GSS dendrogram was, not surprisingly, consistent with the core genome phylogeny, showing the same clade arrangement and relationships for the *Solirubrobacter* (Fig 4c). However, the genome assembly ASM999324v1 reported as *Solirubrobacter* in NCBI, is again located in a clade with *B. subtilis*. The smaller genome of ASM999324v1, along with the phylogenetic inconsistencies, helped us to conclude that it is not a strain of *Solirubrobacter* and was excluded from the rest of the analysis.

*Solirubrobacter* reference genomes harbor an average of 7,423 protein-coding genes, with *S.* sp. URHD having the lowest number and *S. ginsenosidimutans* the highest (6,353 and 8,541, respectively; Fig. 4c). The average GC content was 69.81% and a total of 2,644 proteins were shared among all *Solirubrobacter* species, a strict core genome (Fig. 4d, Fig. S3).

### *Solirubrobacter* diversity and interactions

Because of the large prevalence of *Solirubrobacter* in soil and rhizosphere microbiomes, we started delving into its diversity through sequence retrieval and 16S gene annotation to describe the environmental diversity. We used 107 samples from previous work involving sampling comprehensive Mexican soil diversity and its role in rhizosphere microbiome structuring [14]. *Solirubrobacter* environmental diversity is represented in the phylogeny constructed using the 3,166 *Solirubrobacter* OTUs identified in our samples; we also included the reference genomes showing that they map a few clades from the whole 16S phylogeny (Fig. 4e). *Solirubrobacter*’s diversity shows that a great effort is yet to be carried to capture the diversity of this bacterial genus.

*Solirubrobacter* interactions with other members of the microbial community were evaluated through a network analysis using a 16S co-occurrence network (Fig. 5) from the 107 root-associated and Mexican soil microbiomes. We got a network with 1,748 genera (nodes) and 35,034 edges, representing a large and complex amount of interactions among the bacteria.

**Fig. 5.**
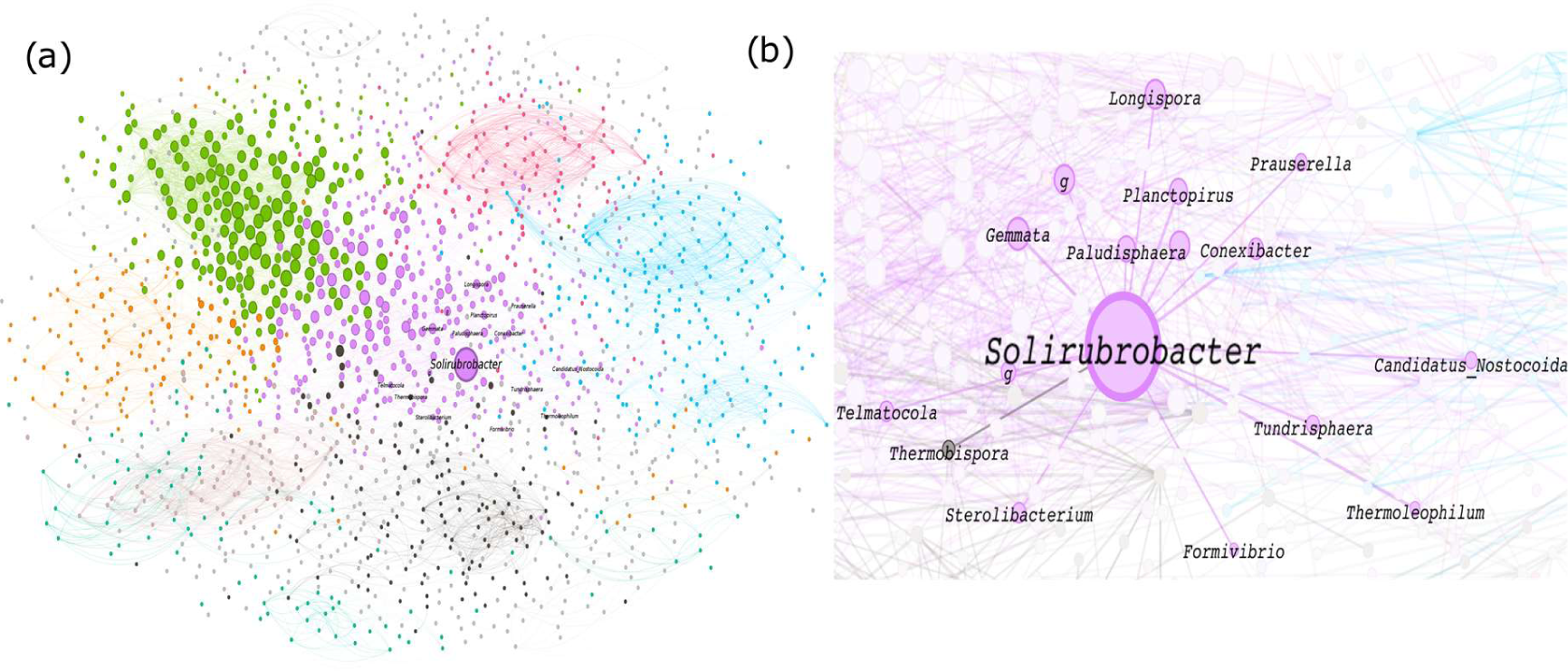
Identifying ecological neighbors for *Solirubrobacter.* (a) Co-occurrence network based in 107 soil and rhizosphere 16S microbiomes highlighting *Solirubrobacter* node. Each node represents a bacterial genus, and each edge a correlation (*r*^2^ > 0.8). Colors represent modularity class. (b) Close-up of the network highlighting the co-occurring genera with *Solirubrobacter*.

The node degree centrality had an average of 40.08 and a range from 0 to 228. *Solirubobacter* had a degree centrality of 97, indicating a larger number of connections to other taxa. Network closeness centrality was approximately 0.020, with *Solirubrobacter* having a larger value of 0.376, suggesting a keystone role. However, *Solirubrobacter* was not a network bridge, as normalized betweenness centrality is low (0.003). To understand community interactions, *Solirubrobacter*’s neighbors according to co-occurrence and higher correlations (*r*^2^ > 0.8) were filtered (Fig. 5b). The *Solirubrobacter* co-occurring genus are: *Gemmata, Longispora, Conexibacter, Planctopirus, Prausurella, Paludisphera, Thermobispora, Sterolibacterium, Thermoleophilum, Tundrisphera, Telmatocola,* and ca. *Nostocoida* (Fig 5b).

### The pangenome as a tool to map *Solirubrobacter* into its naturally occurring niches

We calculated a *Solirubrobacter* pangenome from the genome references, which was composed of 19,645 family proteins, of which 2,644 of them were the genus core genome (Fig. 4d, Fig. S3). A total of 6,917 family proteins were the shell genome and 10,084 the cloud genome, adding up to an accessory genome of 17,001 family proteins. *Solirubrobacter* has an open pangenome, with each new genome adding unique genes (Fig. S4).

Further, we used *Solirubrobacter* pangenome as an anchor to recruit metagenomic reads to assess its environmental distribution in multiple environments. Metagenomic recruitments showed the six-frame translated amino acid identity (AAI) percentage and coverage between *Solirubrobacter* pangenome to metagenomic reads from Mexican soils and rhizospheres (Fig. 6a and 6c, Table S5 and S6). The variation in AAI of translated genes ranges from ∼70-100%, not only in the core genome sequences but throughout the entire pangenome, indicating they mainly belong to the *Solirubrobacter* genus and their ubiquitous presence in Mexican soils and rhizospheres.

**Fig. 6.**
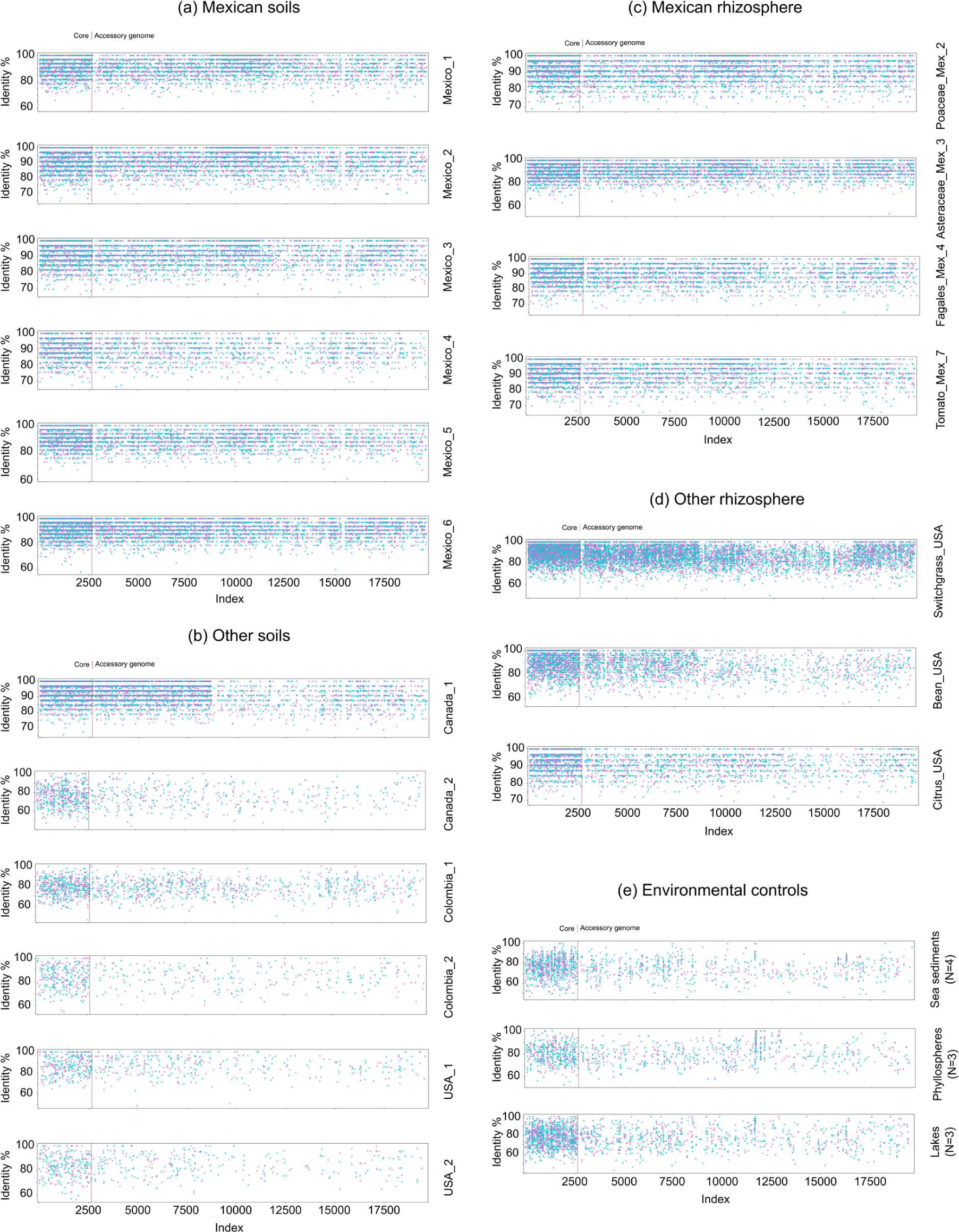
*Solirubrobacter* pangenomic recruitment from metagenomes. Coverage and average amino acid identity (AAI) graphs of (a) Mexican soils, (b) Canadian, Colombian, and American soils, (c) Mexican rhizospheres, (d) American rhizospheres, and (e) control environments. Except for the control environments, each recruitment graph corresponds to a single metagenome of the corresponding environment. Only a representative subset of four Mexican rhizosphere metagenomes with the highest sequencing depth is shown. The recruitment graph for all Mexican rhizospheres is found in Fig. S5. Blue and magenta dots indicate matching six-frame translated sequences in forward or reverse DNA strands. The core genome is to the left of the plots, delimited from the accessory genomes by a vertical brown bar in each graph.

The Mexican rhizosphere encompasses a dataset of tomato *(Solanum lycopersicum*) and wild plants belonging to multiple plant families (*Asteraceae, Fagales, Fabaceae, Lamiaceae*, and *Poaceae*) [14]. Contrary to what is observed for Mexican soils, the metagenomic recruitment with non-Mexican soils was scarce (Fig. 6a and 6b and Table S5 and S7). The percentage of identity recruitments in non-Mexican soils ranged between ∼65 and 100% AAI, with a drastic 50% reduction in recruitments against the accessory genome (Fig. 6b and Table S7). Most of the recruited sequences were from Canadian soil metagenomes, whereas American and Colombian soils presented poor recruitment of *Solirubrobacter* sequences (Fig. 6b and Table S7). On the other hand, non-Mexican rhizospheres presented a higher recruitment and identity percentage throughout the entire pangenome, similar to the profile obtained for Mexican rhizospheres (Fig. 6c and 6d and Table S6 and S8). Nonetheless, rhizospheres from Mexican tomatoes and American citrus and beans presented lower sequence recruitment when compared to the rest of the rhizosphere samples (Fig. 6c and 6d). These recruitment differences suggested that the host genotype may influence the diversity and presence of *Solirubroabcter*.

Regarding lakes, phyllospheres, and sea sediments, the identity percentage ranged between 60 and 90%, with few reads of 100% AAI (Fig. 6e). In these three environments, the sequences recruited were lower than the sequences detected in soils and rhizospheres (Fig. 6). Hence, we can conclude that *Solirubrobacter* is a bacterium associated with rhizospheres and some soils.

Accessory genome sequences with a recruitment coverage ≥ 11 for soils and ≥ 18 for rhizospheres were considered high coverage. This coverage resulted in 13 sequences for Mexican soils (Table S5), including helix_turn_helix, Lux Regulon, MmpL family, and AAA domain of the putative AbiEii Type IV Toxin-Antitoxin system. MmpL proteins have been associated in *Mycobacterium* with translocating siderophores [73]. Concerning rhizospheres, the six high-coverage sequences (Table S6) were the F5/8 type C domain, FGGY family of carbohydrate kinases, histidine kinase HAMP region domain protein, peptidase S8 family involved in serine metabolism, pyridoxal-dependent decarboxylase conserved domain, and short-chain dehydrogenases reductases (SDR) family proteins.

A set of pangenome proteins resulted in gaps in the accessory genome recruitments (gap around the 15,500 index; Fig. 6). These 186 pangenome proteins absent in the metagenomes belong to *S.* sp. URHD, the basal group in both core and GSS phylogenies. Only 14 of these proteins are annotated; the rest are hypothetical proteins. These accessory proteins could be a specialized set of *S.* sp. URHD. Despite being the genome with fewer proteins (6,353; Fig. 4c), *S.* sp. URHD hosted a higher number of unique family proteins (2,124; Fig. 4d). So, it is suggested that *S.* sp. URHD may have undergone an overall genome size reduction, with particular gene expansions and acquisitions to adapt itself to a unique niche. On the contrary, the larger genome sizes of *S. ginsenosidimutans* or *S. taibaiensis* suggest generalist strategies.

### *Solirubrobacter* Environmental Extended Pangenome

We calculated 29,330 protein families comprising the environmental extended pangenome (EEP), meaning 9,906 potentially new environmental *Solirubrobacter* proteins were detected. After COG assignments, 17,189 protein sequences were matched (58.60% of the EEP, Table S9), with the remaining 12,141 as hypothetical proteins. The most abundant COGs were S (unknown; 18.73%), K (transcription; 9.33%), NA(-) (6.74%), E (amino acids; 6.49%), C (energy; 6.19%), T (transduction; 5.71%). We visualized the different COGs per sample as well as the reference core genome and the EEP (Fig. S6). When evaluating the differences among the annotated proteins from the references and our metagenomic annotated proteins, the PCoA ordination revealed two main clusters: an upper cluster containing most of the reference genomes and some metagenomic rhizospheres and a second cluster with almost all soils and the rest of the rhizosphere metagenomes (Fig. S7).

A closer examination of the EPP revealed COGs that increased their abundance compared to the reference pangenome, such as the 23 COGs contained in the lower clusters of the EEP COGs heatmap (Fig. S6). These COGs contain different hydrolases, proteins from the CarA family for carbapenem synthesis (COG EF; amino acids/nucleotide), ethanolamine utilization proteins (COG CQ; energy/secondary metabolism), and phenylacetic acid degradation proteins (COG CI; energy/lipids). Additionally, some COGs were mainly enriched in specific metagenomes. Such is the case of Tomato_Mex_10 COGs MQ (cell wall/secondary metabolism), FI (nucleotides/lipids), DJ (cell cycle/ribosomal), BDLTU (chromatin/cell cycle/replication/transduction/intracellular trafficking), and CI (energy/lipids) (*z* = 3.39 for all of them). The proteins contained in these COGs include methionine biosynthesis protein MetW, ParE protein from the type II toxin-antitoxin system, phosphatidylinositol kinase, and phenylacetic acid degradation proteins. Similarly, samples Mexico_6 and Tomato_Mex_8 contain permease for allantoin in COG FH (nucleotides/coenzymes) (z = 1.82), a plant-derived metabolite that influences microbial community structure in the rhizosphere (Wang et al., 2010). COGs CQ (energy/secondary metabolism) (*z* = 3.39) and EU (amino acids/intracellular trafficking) (*z* = 3.39 for both COGs) from Fagales_Mex_4 contain carboxysome and dienelactone hydrolase proteins, the latter being involved in the catabolism of chlorochatecol.

### *Solirubrobacter* interactions with plants and other microorganisms

We observed a few protein families involved in plant interactions in the reference pangenome. From COG I (lipids), squalene/phytoene synthase was detected, as well as chorismate binding enzyme from COG HQ (coenzymes/secondary metabolism). Conversely, in the EEP, we found protein families involved in chorismate biosynthesis. Regarding motility, 20 protein families are flagellum-related, most found in COG N (motility). Phytase-related protein families were identified in *S. ginsenosidimutans, S. phytolaccae, S.* sp. CPCC, and *S.* sp. URHD. Phytase proteins were identified in *S. ginsenosidimutans, S.* sp. CPCC, and *S. soli*. *S. ginsenosidimutans*, while *S. soli* presented auxin-binding proteins. *S. phytolaccae* contains two unique sequences for phenazine biosynthesis (PhzF and phenazine biosynthesis protein A/B) while sharing three phenazine biosynthesis-like protein sequences with the rest of the species, except for *S.* sp. URHD. *S. phytolaccae* and *S. taibaiensis* contain farnesyl diphosphate synthase (COG MU; cell wall/intracellular trafficking), codified in the *ispA* gene, and involved in carotenoid biosynthesis. Mainly, *S. phytolaccae* contains a Hep/Hag repeat protein domain in COG UW (intracellular trafficking/extracellular structures) (*z* = 1.7638). The Hep/Hag repeat has been associated with bacteria root attachment [74, 75]. However, only 11 sequences from Mexican rhizosphere and soil metagenomes matched a previously reported gene set for plant-microorganism interaction (HMM) (Table S10) [64]. Contrastingly, *S. coelicolor* and *S. griseus* had 142 and 31 positive matches, respectively. The sequences that had a positive match with the HMM were all hypothetical. This matching pattern suggested that *Solirubrobacter* lacks most Actinobacteriota known proteins that foster plant interactions.

## 8. Discussion

### Reference core genome summary of the genus *Solirubrobacter*

The analysis of all the available genomes revealed that strain ASM999324v1 genome did not belong to the genus *Solirubrobacter* (Fig. 4), contradicting the NCBI genome database. The remaining seven *Solirubrobacter* genomes allowed us to construct a 19,645 family protein pangenome. As previously mentioned, *Solirubrobacter* reference pangenome is open (Fig. S4), which is consistent with the fact that *Solirubrobacter* is found in open environments such as soils, in contrast with species with closed pangenomes, which tend to occupy isolated niches [76]. Since the analysis of multiple and independent isolates contributes to the understanding of the global complexity of a bacterial species [76], a more accurate description of this genus can be achieved once the number of available genomes increases.

*Solirubrobacter* ubiquitousness can be explained by its genomic content (Fig. 2). A comprehensive carbohydrate metabolism translates as plasticity when thriving in different environments. Accordingly, COG G (carbohydrate) was among the most abundant in the pangenome and core genome, containing proteins to metabolize simple and complex carbohydrates. Moreover, ROK proteins and cupin domains contribute to *Solirubrobacter*’s metabolic capabilities. The former can act as sugar kinases or as transcriptional regulators involved in carbon and sugar metabolism [77], while the latter is involved in a wide variety of processes, such as acquisition of nutrients, synthesis of antibiotics, and catabolism of different organic compounds [78, 79].

An assortment of sigma factors present in a genome is a proxy of physiological and developmental characteristics, primarily since accessory sigma factors (such as ECF) usually activate the transcription of specific gene sets in response to environmental signals [80]. Selfsame, this bacterium possesses proteins involved in the Lux operon, which regulates the expression of different genes involved in virulence factor expression, exoenzyme secretion, biofilm formation, motility, oxidative stress response, cellulase synthesis, carbon metabolism, RNA processing, sugar transport, and vitamin and biosynthesis of secondary metabolites [81].

Some core proteins allow *Solirubrobacter* to resist different types of environmental stresses, such as oxidative, osmotic, or temperature shock. The presence of antibiotic, chromate, and mycotoxin transporters are defense mechanisms that reduce the intracellular concentration of these hazardous compounds. Glutamate decarboxylase enzyme is involved in the glutamate-dependent acid resistance (GDAR) system, a system described in different bacteria that confers resistance to acidity through scavenging protons by GadA/GadB during GABA formation to elevate internal pH [82, 83]. Finally, the presence of methyltransferases is worth mentioning since they are related to the catabolism of xenobiotic compounds [84]. In agreement with this, *Solirubrobacter* has been associated with degrading different hazardous compounds [85, 86].

### The largest gene families are linked with environmental sensing, defense, and potential motility

Gene families arise from duplicated genes the organism retains because they result in advantageous capacities facing specific environments [87]. *Solirubrobacter* favors gene families involved in environmental sensing, such as the LuxR regulators, and defense, such as antimicrobial compounds (Fig. 3a).

Regarding defense mechanisms, the sequence identified as RNA-mediated transposition proteins was a DDE-type integrase/transposase/recombinase, codified in the *tnsB* gene in *E. coli*. This is a sequence-specific DNA-binding protein required for Tn7 transposition. Tn7 transposon inserts itself at high frequencies and codifies for antibiotic-resistance genes [88].

In Actinobacteriota, LuxR regulators are associated with different domains, and such combinations of domains result in modulation of the expression of different genes, including genes for the biosynthesis of secondary metabolites [81, 89]. Moreover, the specific combination of the LuxR+domain is dependent on the environmental origin of the bacterium, with a positive correlation between the number of LuxR proteins and the association with plants [89]. In Gram-negative bacteria, LuxR is involved in detecting environmental signals, while LuxI contributes to the production of signaling molecules; in Gram-positive bacteria, the presence of LuxR and absence of LuxI could be a technique used by Gram-positive bacteria to interpret Gram-negative quorum-sensing signals [90]. Besides conforming gene families, regulon Lux-related proteins were part of the high-coverage proteins identified in soils. The capacity to sense and interpret environmental stimuli is crucial in highly changing environments, such as soil, especially for a non-motile bacterium like *Solirubrobacter*. This environmental sensing is further sustained by the maintenance of gene families devoted to environmental sensing in the genome of *S.* sp. URHD (Fig. 3a). This URHD strain could be suffering a genome-reduction process.

Regardless of all *Solirubrobacter* isolates being described as non-motile, attention is drawn to the presence of proteins involved in flagellar biosynthesis and flagellin gene families. Despite all *Solirubrobacter* strains harboring flagellin-coding genes, the absence of the cytoplasmic membrane ring (FliF) is noteworthy. This structure plays a pivotal role not only in the flagellum but also in its homologous superstructure, the type III secretion system, and, consequently, the injectisome [91]. Furthermore, the protein MotB, integral to the proton channel and acting as the flagellar motor stator, is conspicuously absent. However, the presence of MotA, the other protein comprising the stator, suggests that all these flagellar genes are not associated with other types of secretion systems [92]. Essential proteins for flagellar biosynthesis vary depending on the species, and the presence of flagellar proteins in non-motile bacteria has been linked to the remains of transport systems [93, 94]. Altogether, these observations lead to the possibility that we may be witnessing vestiges of a flagellum that are no longer in use or that have recruited other proteins in the formation of a novel type of flagellum.

### *Solirubrobacter* is a bacterium associated with soils at a regional scale and with rhizospheres of specific plant hosts

The diversity of *Solirubrobacter*, as revealed by the 16S gene (Fig. 4e), underscores the uncharacterized *Solirubrobacter* species in soils and rhizospheres. This diversity extends to the coding gene level, evidenced by the percentage identity variation of the recruitment graphs for rhizosphere and soil metagenomes (Fig. 6). The quantity and distribution of metagenomic reads recruited to any given genome indicates the abundance of closely related organisms [95]. Conversely, when recruiting metagenomes from lakes, phyllosphere, and sediments, the lower amount and sequence identity of recruited reads suggest a lesser presence of this bacterium in these environments, used as negative environmental controls (Fig. 6e). However, *Solirubrobacter* has been detected as a prevalent endophytic bacteria in the ginseng plant (*Panax notoginseng*) and as one of the two most-abundant genera associated with *Trachymyrmex septentrionalis* ants [96, 97], indicating other environments may also harbor *Solirubrobacter* communities.

The presence of *Solirubrobacter* in soil appears to vary by region: it is found in all Mexican soils, scarcely in Colombian and American soils, while Canadian soils contain *Solirubrobacter’s* core genome but less than 50% of the accessory genome (Fig. 6a and 6b). This variation may indicate the presence of distant *Solirubrobacter* species or closely related bacteria from another possibly unknown genus. As proposed, microbial species distribution is also influenced by local conditions upon immigration to new habitats, much like microorganisms [21, 22]. Therefore, even if *Solirubrobacter* bacteria could disperse from Mexican to foreign soils, varying environmental factors would dictate the persistence or disappearance. Further research is required to explore its presence in other soils and environments, considering abiotic, pedogenic, and regional parameters.

Similarly to soil variations, plant roots appear to affect *Solirubrobacter* abundance and unidentified strains of the genus. Interestingly, *Solirubrobacter* was more abundant and diverse in wild plants’ rhizospheres than in cultivated crops (Fig. 6c and 6d, and Fig. S5). Although not as apparent as soil recruitments, wild plants recruited 9,565 sequences and only 7,910 from tomatoes (Table S6). The higher prevalence of *Solirubrobacter* in wild plants may be explained by changes crops underwent along the domestication process. However, these variations in recruitment may be due to other factors associated with the plant genotype and should be further addressed.

### Environmental Extended Pangenome: Hints for new *Solirubrobacter* species and new metabolic functions

The presence of new *Solirubrobacter* strains was analyzed by comparing the genomic content between reference species and environmental metagenomic samples. The PCoA ordination allowed us to evaluate the difference between the genomic content present in each sample (Fig. S7). The formation of two clusters indicates the presence of two different sets of *Solirubrobacter*: one containing all reference species and some environmental samples and the other one containing species similar between them but different from the references. Hence, it is fair to suggest that these metagenomic samples foster unknown *Solirubroabcter* strains. This idea is further sustained through the percentage identity graphs (Fig. 6) that show enough variation to suggest the presence of new species and through the functional profile of the EEP (Fig. S6), which bears proteins with new functions for the genus.

The construction of a *Solirubrobacter* EEP allowed us not only to identify proteins that potentially belong to this genus but also to suggest a genetic content related to local adaptations while simultaneously broadening our view regarding its metabolic potential. We detected in our metagenomic samples proteins involved in assimilatory sulfate reduction an incomplete route in the reference genomes. Assimilatory sulfate reduction reduces sulfate to sulfide, which can ultimately be used to synthesize cysteine [98]. Sulfur metabolism contributes to soil fertility and the bioavailability of nutrients such as P, Fe, K, and Zn for plants and other microorganisms [99–102]. *Solirubrobacter* from samples Mexico_1, Mexico_4, Asteraceae_Mex_3, and Tomato_Mex_9 seem to reduce sulfate (Fig. S8) partially. The enzyme CisIJ that carries out the last reduction step is present in Mexico_1 and Mexico_4, suggesting that even though *Solirubrobacter*-associated proteins cannot reduce sulfate ultimately, other microorganisms in the environment can carry out the final step, creating a syntrophic process.

### *Solirubrobacter* ecological interactions

In both the reference pangenome and the EEP, permeases for allantoin, chorismate, phytase, and a few other plant-microorganism interaction proteins were identified. Similarly, COG N (motility) was among the COGs with higher representation in the reference pangenome (z = 2.2670). The root-isolated *S. phytolaccae* has a high proportion of the Hep/Hag repeat protein (COG UW; intracellular trafficking/extracellular structures), whose domain has been associated with the binding activity of adhesins, invasins, and agglutinins in soil and plant-associated bacteria [74, 75]. However, *Solirubrobacter* presented ∼30% or less of the plant-microorganism interaction gene sequences [64] that *S. coelicolor* and *S. griseus* bear in their genomes. Moreover, among the high-coverage proteins identified in rhizospheres are proteins that indicate a possible indirect interaction with the plant host. FGGY kinases, a type of high-coverage protein identified in rhizosphere metagenomes, are highly plastic proteins involved in bacterial adaptation to exploit a plethora of carbohydrates using different metabolic pathways [103]. Another high-coverage identified protein in rhizospheres is the pyridoxal-5’-phosphate-dependent enzyme, corresponding to a glutamate decarboxylase. This enzyme is responsible for the first step of GABA synthesis, a compound associated with interactions between plants and bacteria [104].

Despite being associated with the rhizosphere of different plants, the total plant-microorganism interaction genes were relatively low (Table S10) [64]. Hence, even though *Solirubrobacter* has been detected in the rhizosphere of different plants, it appears not to interact with the plant directly. Nevertheless, since we detected a tremendous amount of percentage identity variance between the reference and the metagenomic-recruited proteins, the genetic content of future isolates from rhizospheric environments should be thoroughly analyzed.

Its proximity to the roots could derive from a need to interact with the rest of the bacterial community. Given the absence of biosynthetic pathways for essential amino acids and the lack of numerous enzymes involved in the nitrogen cycle, it is reasonable to surmise that *Solirubrobacter* relies on other microorganisms to fulfill critical metabolic needs. Our network analysis (Fig. 5) revealed that five out of the twelve co-occurring bacteria genera had reported capabilities for nitrate reduction (*Gemmata, Longispora, Conexibacter, Prausurella,* and *Sterolibacterium*) and six of them produce mycelia (*Gemmata, Longispora, Planctopirus, Prausurella, Thermobispora,* and ca. *Nostocoida*) [105–112]. The mycelial structures offer a protective environment for *Solirubrobacter* and other embedded microorganisms. *Planctopirus*, known for its antibiotic production and attachment to eukaryotes, including plants [110], may act as a conduit between *Solirubrobacter* and the plant host. Notably, the antibiotic-producing and multidrug-resistant genus *Gemmata* [105, 113], previously linked to *Solirubrobacter*, was found alongside *Solirubrobacter* as one of the most abundant genera within the core rhizospheric microbiome of invasive buffelgrass (*Pennisetum ciliare*) in a prior study [16]. This suggests a potentially tighter interaction necessary for root colonization and recruitment of the broader microbial community. Moreover, given *Gemmata’s* apparent lack of conventional iron metabolic pathways [105], it may rely on other microorganisms for iron acquisition —the co-occurrence of *Solirubrobacter* and *Gemmata* in a plant-interaction context warrants further investigation.

Our study pioneers an approach beyond employing genomes assembled from metagenomes (MAGs). Initially, we carried out an inventory of genomic diversity based on available databases and characterized the pangenome. This approach expedited comparative genomics studies, aided by a reference. Further, guided by the pangenomic reference, we identified genes within metagenomic assemblies, expanding the pangenomic repertoire of *Solirubrobacter* and shedding light on the dynamics of the communities they inhabit. The methodology employed in this study facilitated a tri-faceted examination of *Solirubrobacter*: phylogenomics, pangenomics, and metagenomic analysis. These perspectives gave us a broader understanding of this bacterial genus, addressing taxonomic diversity and elucidating the plethora of functions inherent in *Solirubrobacter*. Each analysis augmented the number of gene functions associated with *Solirubrobacter* (Fig. 7), underlining the open nature of *Solirubrobacter*’s pangenome while concurrently indicating that specific environments compel their members to possess particular genetic content. Following the discourse “Everything is everywhere, but the environment selects” concerning *Solirubrobacter*, we probed into it on a vast geographical scale, examining North and South American soil and rhizosphere metagenomes. Based on its occurrence patterns, we propose that *Solirubrobacter* is present in every soil, with the rhizosphere being its preferred habitat. We also scrutinized these large-sized genomes, regarding them as generalists [114], aiming to elucidate the mechanisms underlying their successful distribution. *Solirubrobacter*, a non-sporulating Actinobacteriota in all reported isolates, showcases an advantageous trait for environmental endurance and distribution compared to other bacteria like *Bacillus* [115]. *Solirubrobacter* exists as vegetative active cells in the environment, exhibiting drought resistance and compensating for the lack of spore-related DNA damage protection through expanded repair systems like the *uvrABC* gene families reported herein. Notably, despite being a prevalent bacterium, there are no reports of swimming cells, and the pangenomic evidence hints at a deteriorated flagellum system or possibly a novel variant; this warrants further investigation.

**Fig. 7.**
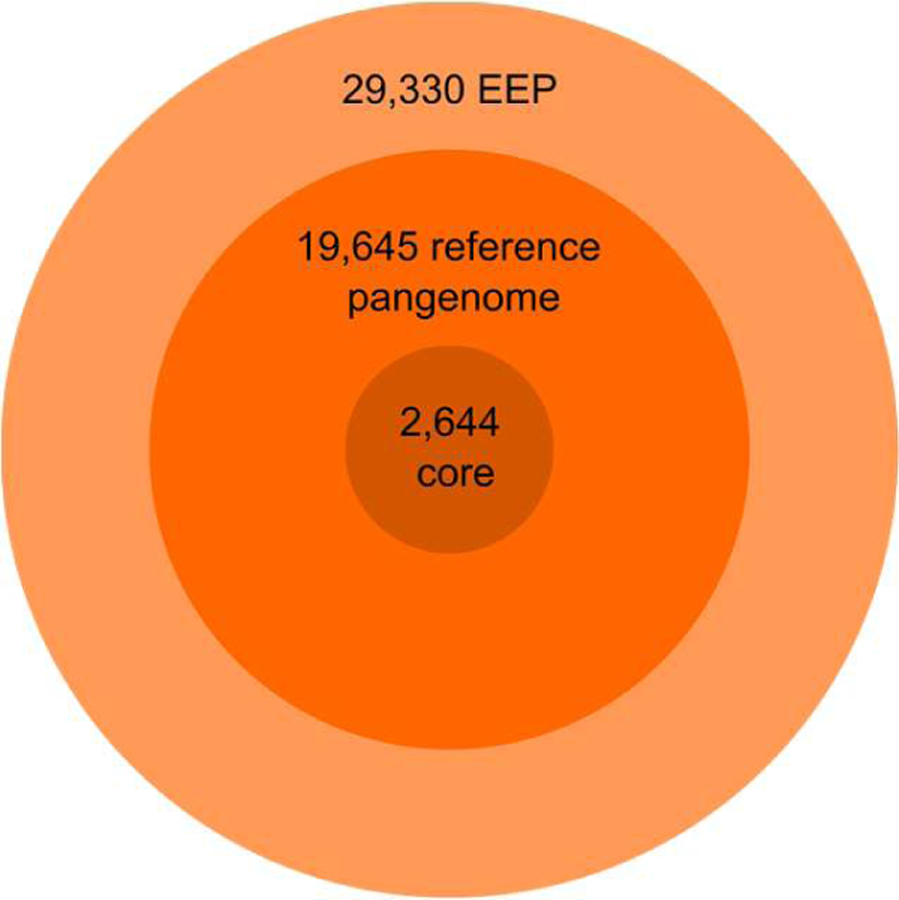
Venn diagram representing protein families in the core genome, the reference pangenome, and the environmental extended pangenome (EEP).

In conclusion, our analysis facilitated the first comparative genomics description of the *Solirubrobacter* genus. The genetic repertoire of *Solirubrobacter* equips it with the capability to metabolize various compounds and endure challenging environments, ranging from defense against microbial adversaries to the degradation of hazardous compounds and adaptation to other environmental abiotic alterations. Intriguingly, its ubiquity is governed not only by its genetic arsenal but also by regional-scale factors. *Solirubrobacter* exhibits a more pronounced prevalence in Mexican soils than in other nations. This pattern insinuates that regional or historical elements may play a role in the dispersal and establishment of *Solirubrobacter*. However, its presence in rhizospheres seems to transcend the geographical origin of the sample, albeit being partially influenced by the host species. These results evoke questions regarding whether bacteria from this genus interact with the plants or confine their interactions to microorganisms associated with the roots. Through this investigation, we augmented the known coding gene functions via the expanded pangenome. Additionally, the pangenomic identity variations observed in pangenome recruitments against metagenomes suggest that the detected *Solirubrobacter* strains possess distinct metabolic capacities, warranting further characterization.

## 9. Author statements

### 9.1 Author contributions

Conceptualization: Angélica Jara-Servín, Luis D. Alcaraz

Data curation: Angélica Jara-Servín, Gerardo Mejia, Miguel F. Romero, Luis D. Alcaraz

Formal analysis: Angélica Jara-Servín, Luis D. Alcaraz

Funding acquisition: Luis D. Alcaraz

Investigation: Angélica Jara-Servín, Luis D. Alcaraz

Methodology: Angélica Jara-Servín, Luis D. Alcaraz

Project Administration: Luis D. Alcaraz

Resources: Mariana Peimbert, Luis D. Alcaraz

Validation: Mariana Peimbert, Luis D. Alcaraz

Visualization: Angélica Jara-Servín, Gerardo Mejia, Luis D. Alcaraz

Writing - original draft: Angélica Jara-Servín, Luis D. Alcaraz

Writing - review & editing: Mariana Peimbert, Luis D. Alcaraz

### 9.2 Conflicts of interest

The authors declare that they do not have a conflict of interest.

### 9.3 Funding information

This work was supported by Universidad Nacional Autónoma de México by the project DGAPA-PAPIIT-UNAM IN206824 to LDA. Conachyt Ph.D. scholarship (CVU 765278) to AJS.

### 9.4 Ethical approval

Not required.

All high-resolution figures can be downloaded from the following URL: https://doi.org/10.6084/m9.figshare.24446521

## Notes

https://github.com/genomica-fciencias-unam/Solirubrobacter

https://doi.org/10.6084/m9.figshare.24446521

https://www.ncbi.nlm.nih.gov/sra/PRJNA603586

https://www.ncbi.nlm.nih.gov/sra/PRJNA603590

https://www.ncbi.nlm.nih.gov/bioproject/603603

### Competing Interest Statement

The authors have declared no competing interest.

