## Supplementary Material for "Unraveling the Genomic and Environmental Diversity of the Ubiquitous *Solirubrobacter*"

Supplementary Figures and Tables are available in the following URL:

https://doi.org/10.6084/m9.figshare.24446521

**Supplementary Figure 1.** Detected aminoacid metabolic routes.
**Supplementary Figure 2.** Predicted structure of the Brevinin-1JDa protein. (A) Structure of Brevinin-1JDa. (B) Predicted structure of protein family number 528 that showed a Z value. Analysis was carried out using AlphaFold-Colab (Mirdita et al., 2022). Protein structure prediction was run with default parameters and a PBD model with the highest pLDDT score (pLDDT=65.1). Structure similarity of the predicted protein was determined using the Advanced Search Query Builder Tool from RCSB Protein Data Bank (Berman et al., 2000) (https://www.rcsb.org/search/advanced/structure). Strict search, assemblies, and Computed Structure Models (CSM) parameters were selected. A structural alignment was carried out using PyMOL v. 2.5 (The PyMOL Molecular Graphics System, Schrödinger, LLC.) for protein structure visualization.
**Supplementary Figure 3.** Shared protein families between *Solirubrobacter* references were visualized in an UpSet plot. The histogram shows the number of shared elements for each intersection set, ordered in a decreasing manner.
**Supplementary Figure 4.** Total number of new and conserved genes through sequentially adding genomes to the pangenome.
**Supplementary Figure 5.** *Solirubrobacter* recruitment from soil, rhizospheres, and environmental controls metagenomes. Coverage and percentage identity graphs of (a) Mexican soils, (b) Canadian, Colombian, and American soils, (c) all Mexican rhizospheres, (d) American rhizospheres, and (e) environmental controls. Blue and magenta dots indicate the sequence was found in the sense or antisense chain. Core and accessory genomes are delimited by a vertical brown bar in each graph. Translated nucleotide sequences from the pangenome into the six possible reading frames were used as a query to search in *Solirubrobacter* metagenomic reads. Specific sampling locations for each metagenome can be found in Table S1.
**Supplementary Figure 6.** Functional profile of *Solirubrobacter* Environmental Extended Pangenome. Heatmap showing normalized Cluster of Orthologous Groups (COGs) frequencies within each group are shown as a two-way clustering. Circles refer to reference strains, squares to metagenomic samples, and stars to the reference and environmental extended pangenomes.

**Supplementary Figure 7.** *Solirubrobacter* diversity in environmental metagenomes. Principal Coordinates Analysis (PCoA) unconstrained ordination using Bray-Curtis dissimilarity. Statistical significance was evaluated using the ANOSIM test for environmental origin ($p = 0.0004$) and the formed clusters ($p = 0.0004$).

**Supplementary Figure 8.** *Solirubrobacter* enzymes are involved in sulfur metabolism. The known enzymes involved in the assimilatory sulfate reduction route are shown at the top. The presence of these enzymes is highlighted in blue when present in the *Solirubrobacter* metagenomic proteins or the *Solirubrobacter* reference genomes. Enzymes highlighted in green indicate the presence of the protein in the rest of the proteins of the metagenomes, meaning they are present in other microorganisms but not in *Solirubrobacter*.
